# Interactive online brain shape visualization

**DOI:** 10.1101/067678

**Authors:** Anisha Keshavan, Arno Klein, Ben Cipollini

## Abstract

Urbanization presents unique environmental challenges to human commensal species. The Afrotropical Anopheles gambiae complex contains a number of synanthropic mosquito species that are major vectors of malaria. To examine ongoing cryptic diversification within the complex, we performed reduced representation sequencing on 941 mosquitoes collected across four ecogeographic zones in Cameroon. We find evidence for clear subdivision within An. coluzzii and An. gambiae s.s. - the two most significant malaria vectors in the region. Importantly, in both species rural and urban populations of mosquitoes were genetically differentiated. Genome scans of cryptic subgroups reveal pervasive signatures of selection centered on genes involved in xenobiotic resistance. Notably, a selective sweep containing eight detoxification enzymes is unique to urban mosquitoes that exploit polluted breeding sites. Overall, our study reveals that anthropogenic environmental modification is driving population differentiation and local adaptation in African malaria mosquitoes with potentially significant consequences for malaria epidemiology.

## 1 Introduction

Our goal for the hackathon was to create an interactive Web browser application to visualize humanbrain image data processed by the Mindboggle software package [1].

The Mindboggle project was initiated to improve the labeling as well as morphometry of brain imaging data, and to promote open science by making all data, software, and documentation freely and openly available. An interface for interactive visualization is essential for assessing issues in brain image processing and analysis, including surface reconstruction, labeling, and morphometry. Mindboggle processes human brain cortical surface meshes in the VTK format, and generates label and shape information for each anatomical region, where labels follow the Desikan-Killiany-Tourville protocol [2].

## 2 Approach

Over the course of two afternoons at the Human Brain Mapping 2015 conference’s hackathon, we evaluated several JavaScript libraries for creating browser-based WebGL visualizations of brain surfaces, including three.js, XTK, and BrainBrowser. Three.js was chosen for ease of use and degree of active development and community support. To accompany these surface visualizations with graphical plots, we chose the d3 JavaScript library for its flexibility and widespread use.

## 3 Results

We completed an initial version of our browser-based interactive visualization tool; a left hemisphere of a human brain is available at http://roygbiv.mindboggle.info. Click and drag to rotate this brain, scroll to zoom in and out, and click on any region of the brain while pressing the shift key to produce an accompanying plot of shape measures for that region (fig. 2). This will render all otherregions transparent. Figure 3 shows the distributions of travel depth, geodesic depth, mean curvature, freesurfer curvature, and freesurfer cortical thickness for the selected region. Shift-click outside the brain to return opacity to all regions.

**Figure 1.**
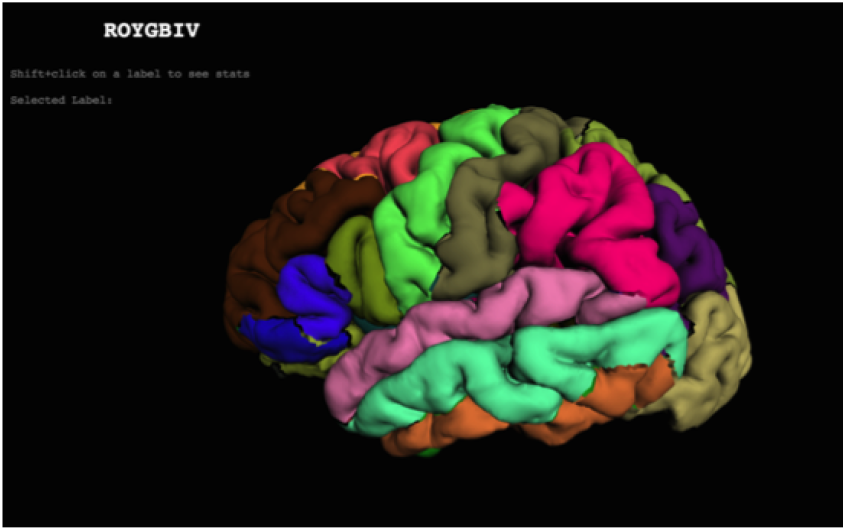
Example visualization.

**Figure 2.**
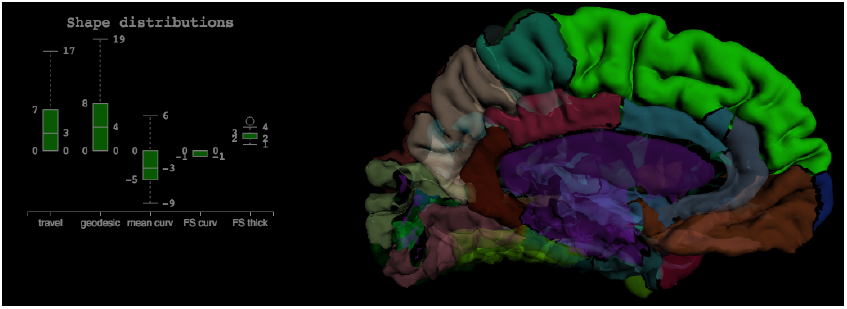
Example of a selected region and the accompanying boxplot.

**Figure 3.**
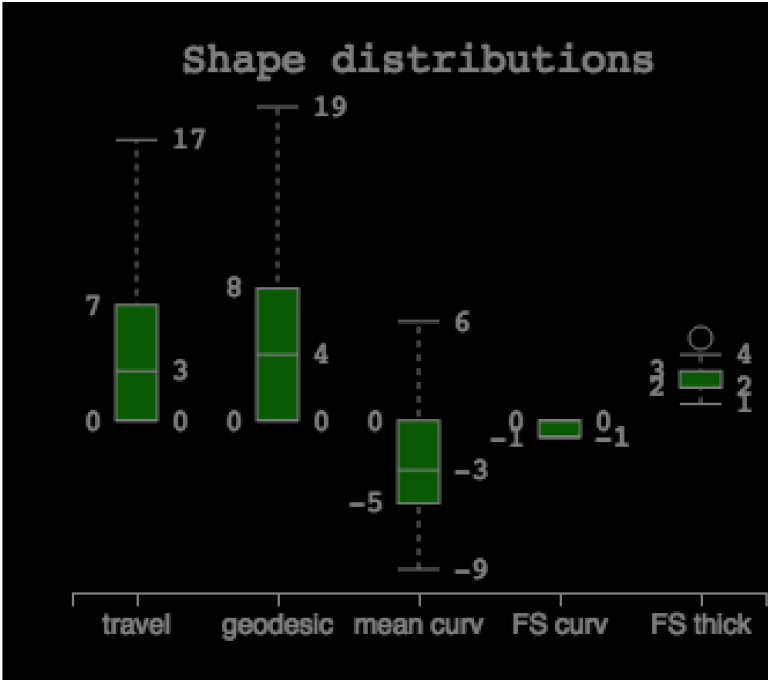
Example boxplot of a selected region that shows the distributions of shape features.

After the hackathon, we refactored the code to use an object-based approach. This allows multiple brains to be shown simultaneously. This approach was used to create a master-slave interaction: selection of a ROI in one hemisphere loads data for display on a second hemisphere. This approach was used in a dynamic poster presented at Society for Neuroscience in 2015 [3].

**Figure 4.**
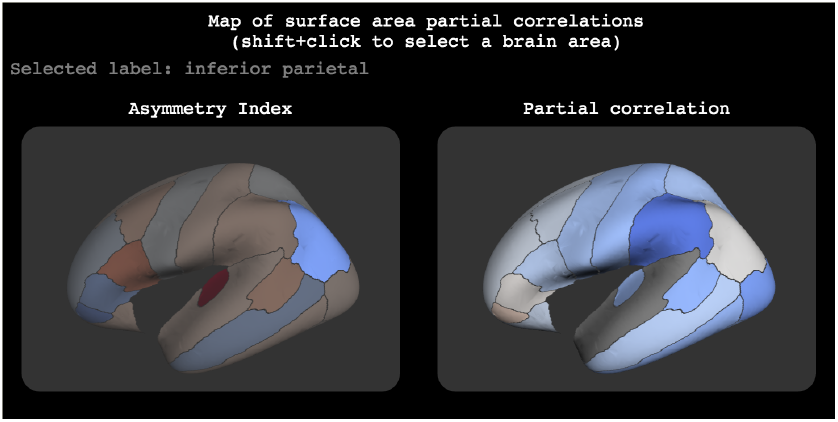
Example master/slave visualization.

## 4 Conclusions

We have received very positive feedback for our efforts at the hackathon, and have since received several requests and encouragement to build this visualization out to accommodate other data besides shape information and to enable the visual evaluation of thousands of brains. We hope to continue this work with the help of others! To contribute to this project, please send pull requests to https://github.com/binarybottle/roygbiv.

## Availability of Supporting Data

More information about this project can be found at: https://github.com/binarybottle/roygbiv. Further data and files supporting this project are hosted in the *GigaScience* repository REFXXX.

## Author’s contributions

AK and AK performed the project and wrote the report. BC extended this work and is actively maintaining ROYGBIV.

## Acknowledgements

The authors would like to thank the organizers and attendees of the 2015 OHBM Hackathon. This project is supported in part by a grant from the NSF (award 1429999).

## References

1. Klein, A., Hirsch, J.: Mindboggle: a scatterbrained approach to automate brain labeling. Neuroimage 24(2), 261–280 (2005)

2. Klein, A., Tourville, J.: 101 labeled brain images and a consistent human cortical labeling protocol. Frontiers in Neuroscience 6(171) (2012). doi:10.3389/fnins.2012.00171

3. Cipollini, B., Bartsch, H., Cottrell, G.: Exploring the anatomy and genetics of cortical asymmetries in surface area and thickness. In: 45th Annual Meeting of the Society for Neuroscience, Chicago (2015)

